# Glucocorticoid Programming of Erythrocyte Hypoxic Memory Enables Rapid High-Altitude Acclimatization

**DOI:** 10.64898/2026.05.26.728051

**Authors:** Tianshen Chou, Yujin Zhang, Xiangpei Yu, Jiayue Gao, Changhan Chen, Wuping Liu, Juan Liu, Zhiyu Yang, Rodney E. Kellems, Lingling Zhu, Yang Xia

**Affiliations:** Department of Otolaryngology Head and Neck Surgery, Xiangya Hospital, Central South University, Changsha, Hunan, China; National Medical Metabolomics International Collaborative Research Center, Xiangya Hospital, Central South University, Changsha, Hunan, China; National Clinical Research Center for Geriatric Disease (Xiangya Hospital), Changsha, Hunan, China; FuRong Laboratory, Changsha, Hunan, China; Department of Brain Protection and Plasticity, Beijing Institute of Basic Medical Sciences, Beijing, China; Department of Biochemistry and Molecular Biology, The University of Texas McGovern Medical School at Houston, Houston, Texas

**Keywords:** Intermittent Hypoxia Training (IHT), High Altitude, Erythrocytes, Glucocorticoid

## Abstract

**Background:** Rapid ascent to high altitude causes acute mountain sickness (AMS) and life-threatening pulmonary/cerebral edema, yet no prophylaxis enables immediate acclimatization. Intermittent hypoxia training (IHT) establishes a “hypoxic memory” that accelerates adaptation, to high altitude, but its cellular and molecular basis remains undefined, precluding effective pharmacological strategies.

**Methods:** A human cohort of 18 sea-level inhibitants was equally divided into two groups, one group received IHT prior to ascent to 3,500 meters, the other group did not. Multi-omics profiling of erythrocytes and plasma, along with isotopic glucose tracing, was employed to examine the metabolic effects of IHT upon high altitude acclimatization. Preclinical studies with genetically engineered mice were used to further define the molecular and metabolic basis of IHT-induced hypoxic memory allowing rapid acclimatization to high altitude.

**Results:** Metabolomics revealed glucocorticoids as previously unrecognized endogenous erythroid hypoxic memory orchestrators induced by IHT that negatively correlated with AMS severity. Lipidomics and isotopic glucose tracing demonstrated that glucocorticoid signaling via the glucocorticoid receptor (GR) coordinately enhanced glucose metabolism and activated sphingosine kinase-1 (SPHK1)-driven sphingosine-1-phosphate (S1P) synthesis, pre-conditioning erythrocyte oxygen unloading and antioxidant capacity. Glucocorticoid supplementation enhanced erythrocyte SPHK1 activation and oxygen delivery, counteracting multi-tissue hypoxia and pulmonary and renal neutrophil infiltration. Conversely, erythrocyte-specific *Sphk1* ablation abolished glucocorticoid-induced S1P production causing severe tissue hypoxia and exaggerated pulmonary neutrophil infiltration.

**Conclusions:** We establish a new function of glucocorticoids in erythrocyte metabolic plasticity to enhance oxygen delivery as a hypoxic memory mechanism for rapid adaptation to high altitude. This previously unrecognized GR-mediated reprograming of glucose and sphingolipid metabolism offers a transformative precision pharmacologic strategy for high altitude preconditioning, high altitude emergencies and hypoxia-driven diseases.

## INTRODUCTION

Rapid ascent to high-altitude environments poses significant health risks due to hypobaric hypoxia, which can precipitate acute mountain sickness (AMS) and potentially fatal complications such as high-altitude pulmonary edema and high-altitude cerebral edema^1–6^. These conditions represent a spectrum of hypoxia-induced pathologies that may progress from mild symptoms to life-threatening multi-organ dysfunction within hours, especially in individuals lacking innate or acquired hypoxic tolerance^7, 8^. While current preventive strategies, including staged acclimatization and pharmacological prophylaxis, offer some protection, they remain only partially effective, underscoring the need for more robust pre-acclimatization approaches^9^. Intermittent hypoxia training (IHT), a non-pharmacological intervention involving repeated exposure to normobaric hypoxia, is widely used to induce hypoxic preconditioning prior to ascent^10–12^. Nevertheless, the precise mechanisms that enable IHT to confer rapid adaptation remain poorly understood.

IHT is well known to enhance hypoxic tolerance by activating endogenous adaptive pathways that produce multisystem protective effects^11, 13–15^. Although prior research has largely focused on its benefits for circulatory, respiratory, and neurological functions, the role of erythrocytes, the primary carriers of oxygen, in IHT-induced adaptation has been largely overlooked^16–20^. Emerging evidence indicates that mature erythrocytes possess active regulatory and stress-responsive capacities that extend well beyond their classical role as passive gas transporters^21–26^. For instance, erythrocytes can sense hypoxia through adenosine-activated signaling pathways such as ADORA2B-AMPK and dynamically modulate oxygen offloading by adjusting 2,3-bisphosphoglycerate (2,3-BPG) levels, thereby fine-tuning tissue oxygen delivery^21, 24^. Despite these advances, it remains unclear whether IHT promotes rapid adaptation to high-altitude hypoxia via such erythrocyte-based “hypoxic memory” mechanisms. The specific metabolic and molecular foundations of the IHT preconditioning process are still not well defined. Given that metabolic reprogramming constitutes a fundamental cellular strategy for adapting to environmental stress, a systematic characterization of the erythrocyte metabolic profile following IHT and after high-altitude exposure is essential to elucidate the underlying adaptive machinery.

To identify the potential “hypoxia memory” apparatus, we employed multi-omics profiling and isotopic glucose tracing in human erythrocytes, complemented by murine genetic models and preclinical studies. Here we show that an IHT-dependent glucocorticoid-driven “GC-GR-SPHK1-S1P” signaling axis in erythrocytes orchestrates a dual metabolic-proteostatic program encoding hypoxic memory. Glucocorticoids signal via GR to coordinately enhance glycolysis and activate SPHK1-driven S1P synthesis, generating a glucose-sphingolipid metabolic switch that pre-conditions oxygen unloading capacity for rapid acclimation. These findings establish erythrocyte glucocorticoid pre-conditioning mechanistically distinct from its standard systemic anti-inflammatory suppression prophylaxis by directly programming erythrocyte oxygen delivery and proteostatic resilience as a translatable strategy for high-altitude hypoxia and hypoxia-related cardiovascular diseases.

## METHODS

### Human participants

This study enrolled 18 healthy young volunteers (12 males, 6 females, mean age 22.13 ± 1.86 years). All participants provided written informed consent. Exclusion criteria were: (1) birth or residence >1500 m, (2) travel to elevations >1000 m within the preceding 3 months, (3) current use of prescription medication, (4) smoking, (5) personal or family history of migraine, and (6) known hematologic or cardiovascular disorders. The protocol was approved by Ethics Committee of Beijing Institute of Basic Medical Sciences (approval number AF/SC-08/02.328) and conducted in accordance with the Declaration of Helsinki.

### Mouse models

Erythrocyte-specific *Sphk1*-knockout (*eSphk1^−/−^*) mice were generated by crossing mice homozygous for a floxed *Sphk1* allele (a kind gift from Dr. Richard L. Proia, NIDDK) with mice expressing Cre recombinase under the control of erythropoietin receptor (EpoR) gene regulatory elements (a kind gift from Dr. Stuart H. Orkin, Boston Children’s Hospital). Both lines have been previously described and were maintained on a C57BL/6 background^27, 28^. All animal procedures were performed in accordance with the US National Institutes of Health guidelines and were approved by the Experimental Animal Ethics Committee of Central South University (Approval No. CSU-2023-0336). Mice were housed under specific pathogen-free conditions at 22 °C with 50% humidity and a 12-h light/dark cycle (lights on from 7:00 AM to 7:00 PM), with ad libitum access to food and water.

For glucocorticoid (GC) treatment, cortisol and corticosterone (both from Aladdin) were first dissolved in DMSO to prepare stock solutions, which were then diluted in PBS to obtain working solutions. Mice received intraperitoneal injections of GCs at a dose of 0.5 mg/kg daily until the day before hypoxic chamber exposure.

### Human study design and IHT protocol

Participants were randomly assigned to an intermittent hypoxia training (IHT) group (n = 9) or a control group (n = 9). The IHT group completed a 7-day normobaric hypoxic training protocol at sea level using a GO2 Altitude Hypoxicator. The daily regimen comprised eight 1-hour sessions, each consisting of alternating 5-minute periods of mild hypoxia (targeting peripheral oxygen saturation, SpO₂, of 80%–85%, equivalent to an inspired O₂ fraction of ∼11%–12%) with 3-minute intervals of breathing normoxic air. Heart rate, SpO₂, and inspired oxygen concentration were continuously monitored. The control group received no intervention. Both groups were subsequently exposed to a simulated high altitude of 3,500 m for 2 days.

### Human sample collection and physiological assessment

Physiological assessments and venous blood collection were performed at four time points for the IHT group: before the training period (Before IHT), immediately after the 7-day training period while still at sea level (After IHT), and on the first (HA-D1) and second (HA-D2) day at high altitude. The control group was assessed at sea-level, HA-D1, and HA-D2 time points. Assessments included the Lake Louise Score (LLS) for AMS severity, peripheral oxygen saturation (SpO₂), heart rate, and a complete blood count (CBC).

### Blood sample processing

Venous blood was drawn into EDTA-coated tubes at approximately noon on each sampling day. Samples were immediately centrifuged at 2,000 g for 5 min at 4 °C. Plasma and the buffy coat were carefully aspirated. The pelleted erythrocytes were washed once with five volumes of ice-cold phosphate-buffered saline (PBS) and aliquoted for subsequent analysis. All samples were snap-frozen in liquid nitrogen and stored at −80 °C until use.

### Untargeted metabolomics profiling

Erythrocyte (20 μL) or plasma (20 μL) samples were extracted in ice-cold methanol:acetonitrile:water (5:3:2, v/v) as described^29^. After vortexing (1 h, 4 °C) and centrifugation (10,000 g, 10 min, 4 °C), supernatants were analyzed on a Vanquish UHPLC system coupled to a Q Exactive mass spectrometer (Thermo Fisher Scientific). Metabolites were separated on a Kinetex C18 column (2.1 × 15 mm, 1.7 μm) using a 5-min gradient. MS data were acquired in full-scan mode (65–900 m/z, resolution 70,000) in positive and negative ionization. Raw data files were converted to mzXML format and processed using the Maven software suite (Princeton University) for peak detection, alignment, and integration. Quality control samples were interspersed every ten injections.

### Untargeted lipidomics profiling

Erythrocyte lipids were extracted with cold isopropanol. After incubation (−20 °C, 1 h) and centrifugation (11,000 g, 10 min, 4℃), supernatants were analyzed by LC-MS/MS on an Orbitrap Exploris 240 system (Thermo Fisher Scientific). Lipids were separated on a C18 column with a gradient of (A) acetonitrile:water (60:40) containing 0.05% formic acid/10 mM ammonium acetate and (B) isopropanol:acetonitrile (90:10) with the same additives. MS data were acquired in positive/negative modes (resolution 60,000 for MS1, 30,000 for MS/MS). Lipid identification and quantification were performed using LipidSearch software (version 4.2, Thermo Fisher Scientific).

### ¹³C-Glucose metabolic tracer analysis

Freshly isolated human erythrocytes were resuspended in glucose-free RPMI 1640 medium supplemented with 6 mM ¹³C-D-glucose (Cambridge Isotope Laboratories). Cells were incubated for 30 min at 37°C under normoxic or hypoxic conditions as indicated. The reaction was quenched with ice-cold methanol:acetonitrile:water (5:3:2). After extraction and centrifugation, the supernatant was analyzed by the same UHPLC-MS platform used for untargeted metabolomics. Isotopologue distributions of glycolytic and pentose phosphate pathway intermediates were analyzed using the Maven software^30^.

### Targeted quantification of glucocorticoids by LC-MS/MS

Glucocorticoids were measured as described^31^. Briefly, 50 μL of erythrocyte or plasma extract was mixed with acetonitrile/MTBE (1:9), vortexed, and centrifuged. The supernatant was dried under N₂, reconstituted in 60% methanol, and analyzed on an ACQUITY UPLC I-Class system coupled to a SCIEX Triple Quad 6500^+^ mass spectrometer. Multiple reaction monitoring (MRM) was used for cortisol and corticosterone quantification.

### Erythrocyte functional assays in vitro

#### Oxygen dissociation curve and P50 measurement

Oxygen equilibrium curves were obtained using a Hemox Analyzer (TCS Scientific Corp.) as described^32, 33^. Briefly, 10 µL of whole blood was mixed with Hemox Buffer, antifoaming reagent, and bovine serum albumin. The sample was equilibrated with varying O₂ tensions at 37 °C, and hemoglobin oxygen saturation was measured spectrophotometrically. The P50 value (partial pressure of O₂ at 50% hemoglobin saturation) was derived from the curve.

#### Intracellular ROS measurement

Erythrocytes were incubated with 2 μM 2′,7′-dichlorodihydrofluorescein diacetate (H₂DCFDA) in PBS for 45 min at 37 °C in the dark. Cells were washed three times, and fluorescence intensity was measured using a Synergy HTX multi-mode microplate reader (BioTek) with excitation/emission at 485/528 nm.

#### SPHK1 activity assay

Erythrocyte SPHK1 activity was measured using a fluorescence-based assay with NBD-labeled sphingosine (NBD-Sph) as the substrate^34^. Briefly, erythrocytes were lysed, and the supernatant was collected as the enzyme source. The enzymatic reaction was performed in a buffer containing 50 mM HEPES (pH 7.4), 15 mM MgCl₂, 0.05% Triton X-100, 10 µM NBD-Sph, and 1 mM ATP. After incubation at 37 °C for 30 min, the reaction was terminated, and the fluorescent product NBD-S1P was extracted and quantified by measuring fluorescence (excitation/emission = 489/535 nm) on a microplate reader. A standard curve of NBD-S1P was used for absolute quantification. Enzyme activity was normalized to the total protein concentration of each sample.

### In vivo tissue hypoxia detection

Tissue hypoxia was assessed using the Hypoxyprobe™-1 (pimonidazole HCl) kit (Hypoxyprobe, Inc.). Mice received an intraperitoneal injection of pimonidazole (60 mg per kg body weight). Thirty minutes post-injection, mice were euthanized, and target tissues (kidney, brain, liver) were harvested, fixed in 4% paraformaldehyde, and embedded in paraffin. Tissue sections (5 μm) were deparaffinized, rehydrated, and subjected to antigen retrieval. Sections were incubated overnight at 4°C with a rabbit anti-pimonidazole primary antibody (1:200 dilution), followed by a 1-h incubation with an Alexa Fluor 488-conjugated donkey anti-rabbit IgG secondary antibody (1:1000 dilution) at room temperature. Nuclei were counterstained with DAPI. Fluorescence images were acquired using a confocal microscope, and the integrated fluorescence density of Hypoxyprobe staining from at least 20 random fields per sample was quantified using Image-Pro Plus software (Media Cybernetics).

### Statistical analysis

Data are presented as mean ± standard deviation (SD). Statistical analyses were performed using R (version 4.4.1) and GraphPad Prism (version 9.0). Normality was assessed using the Shapiro-Wilk test. For comparisons between two groups, unpaired or paired two-tailed Student’s t-tests were used as appropriate. For multiple comparisons, one-way or two-way analysis of variance (ANOVA) was applied, followed by Tukey’s post-hoc test for pairwise comparisons. The Pearson correlation coefficient was used to assess linear relationships. Pathway enrichment analysis of metabolomics data was performed using the clusterProfiler R package. A P-value of < 0.05 was considered statistically significant (*P < 0.05, **P < 0.01,***P < 0.001).

## RESULTS

### Glucocorticoids Are Identified as Key “Hypoxia Memory Mediators” Induced by IHT

To investigate the molecular basis by which intermittent hypoxia training (IHT) enhances resistance to acute mountain sickness (AMS) in sea-level residents, we conducted a human high-altitude study comparing individuals who had undergone IHT, with matched controls who had not **(Fig. 1A)**. Briefly, 18 volunteers were randomly assigned to an IHT group or a control group. The IHT group completed a 7-day IHT protocol at sea level, with blood samples collected before and after training. The control group received no intervention and was sampled at sea level. Both groups were subsequently exposed to high altitude (HA, 3,500 m) for two days, with samples collected on HA day 1 (HA-D1) and HA day 2 (HA-D2). Lake Louise Scores (LLS, an assessment of acute mountain sickness) were determined at sea level (pre/post IHT for the IHT group) and on HA-D1 and HA-D2 for all participants **(Fig. 1A).** Complete blood count (CBC) parameters for both cohorts are summarized in **Tables 1** and **2**.

**Figure 1.**
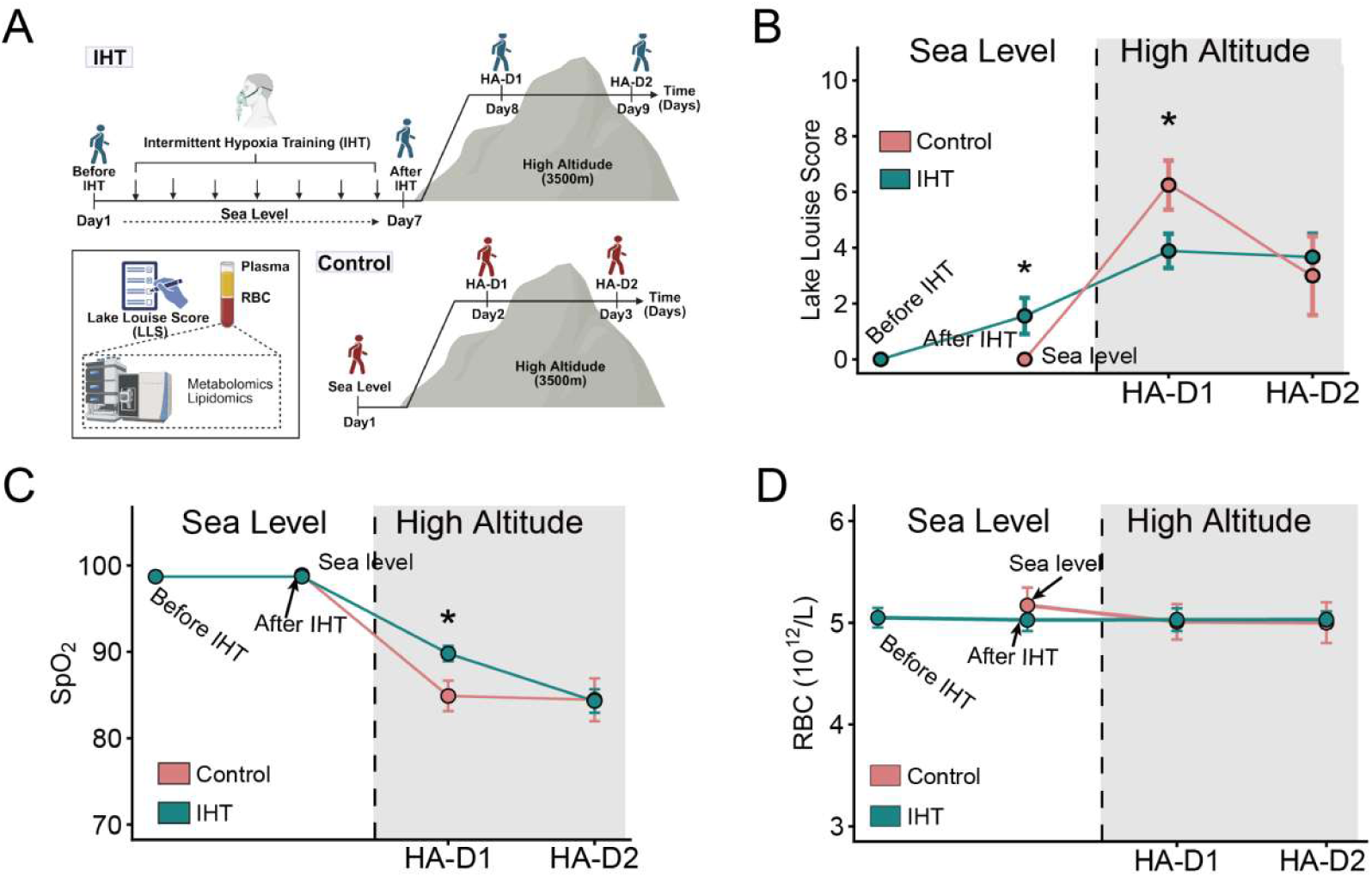
Pre-acclimatization with Intermittent Hypoxic Training attenuates symptoms of Acute Mountain Sickness and improves systemic oxygenation upon high-altitude exposure. **(A)** Schematic of the study design. Participants were randomly assigned to an Intermittent Hypoxic Training (IHT) group (upper panel, blue timeline) or a normoxic Control group (lower panel, red timeline). The IHT group underwent a 7-day (D1-D7) normobaric hypoxic training regimen prior to ascent. Both groups were subsequently assessed at high altitude (HA; 3,500 m) for 2 days (D8-D9 for the IHT group; D2-D3 for the Control group). Venous blood was collected at the time points for subsequent plasma and red blood cell (RBC) metabolomic and lipidomic analyses. **(B)** Lake Louise Score (LLS). a composite clinical score for acute mountain sickness severity, was obtained for all participants at the times indicated. (C) Peripheral oxygen saturation (SpO₂). **(D)** Red blood cell (RBC) count. Data are presented as mean ± SEM, *p < 0.05 (ANOVA followed by Šidák’s post hoc test). P values refer to the differences between IHT and control groups on the day indicated.

**Table1.**
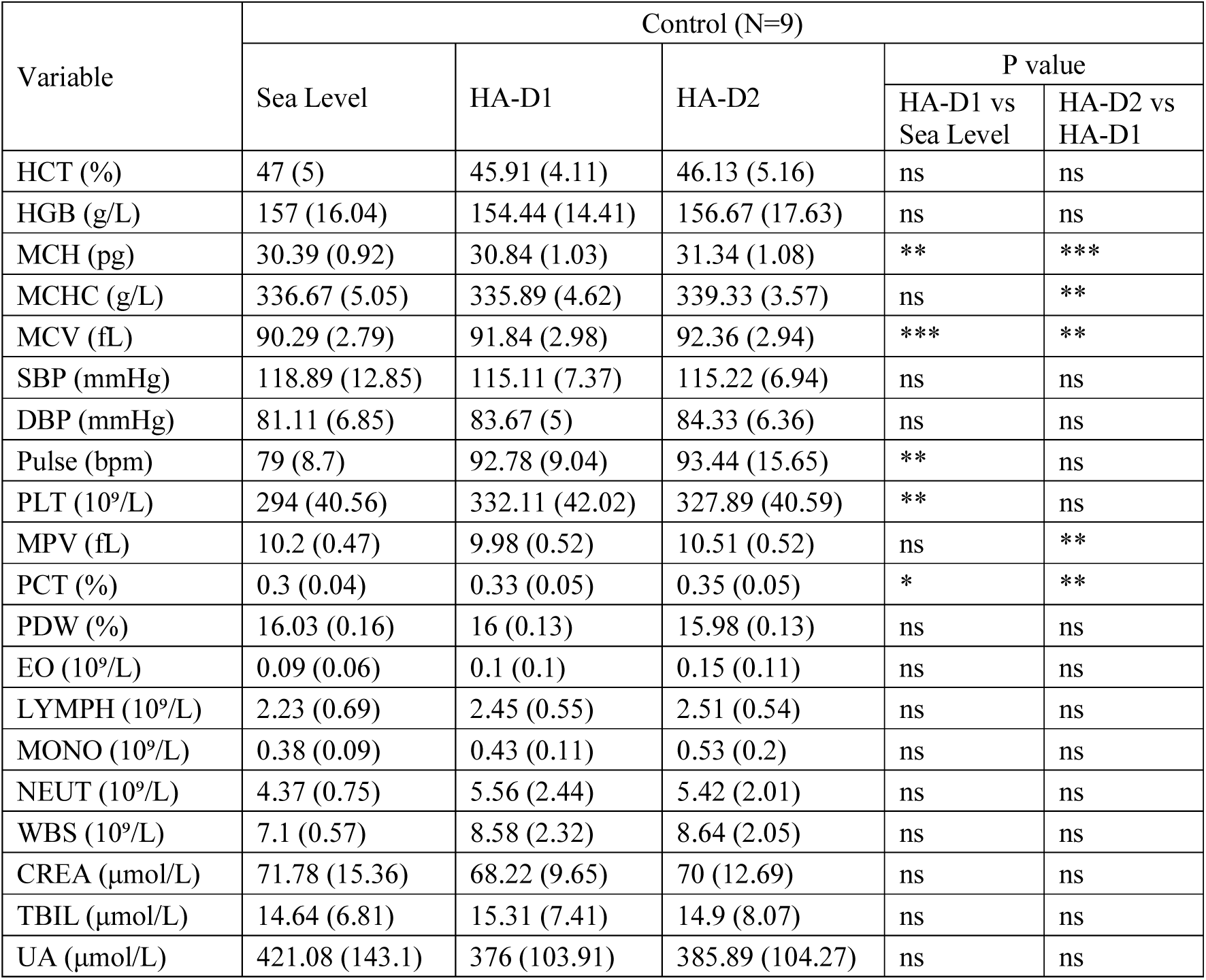
Demographic and clinical information of the control group.

| Variable | Control (N=9) |  |  |  |  |
| --- | --- | --- | --- | --- | --- |
|  | Sea Level | HA-D1 | HA-D2 | P value |  |
|  |  |  |  | HA-D1 vs<br>Sea Level | HA-D2 vs<br>HA-D1 |
| HCT (%) | 47 (5) | 45.91 (4.11) | 46.13 (5.16) | ns | ns |
| HGB (g/L) | 157 (16.04) | 154.44 (14.41) | 156.67 (17.63) | ns | ns |
| MCH (pg) | 30.39 (0.92) | 30.84 (1.03) | 31.34 (1.08) | ** | *** |
| MCHC (g/L) | 336.67 (5.05) | 335.89 (4.62) | 339.33 (3.57) | ns | ** |
| MCV (fL) | 90.29 (2.79) | 91.84 (2.98) | 92.36 (2.94) | *** | ** |
| SBP (mmHg) | 118.89 (12.85) | 115.11 (7.37) | 115.22 (6.94) | ns | ns |
| DBP (mmHg) | 81.11 (6.85) | 83.67 (5) | 84.33 (6.36) | ns | ns |
| Pulse (bpm) | 79 (8.7) | 92.78 (9.04) | 93.44 (15.65) | ** | ns |
| PLT (10<sup>9</sup>/L) | 294 (40.56) | 332.11 (42.02) | 327.89 (40.59) | ** | ns |
| MPV (fL) | 10.2 (0.47) | 9.98 (0.52) | 10.51 (0.52) | ns | ** |
| PCT (%) | 0.3 (0.04) | 0.33 (0.05) | 0.35 (0.05) | * | ** |
| PDW (%) | 16.03 (0.16) | 16 (0.13) | 15.98 (0.13) | ns | ns |
| EO (10<sup>9</sup>/L) | 0.09 (0.06) | 0.1 (0.1) | 0.15 (0.11) | ns | ns |
| LYMPH (10<sup>9</sup>/L) | 2.23 (0.69) | 2.45 (0.55) | 2.51 (0.54) | ns | ns |
| MONO (10<sup>9</sup>/L) | 0.38 (0.09) | 0.43 (0.11) | 0.53 (0.2) | ns | ns |
| NEUT (10<sup>9</sup>/L) | 4.37 (0.75) | 5.56 (2.44) | 5.42 (2.01) | ns | ns |
| WBS (10<sup>9</sup>/L) | 7.1 (0.57) | 8.58 (2.32) | 8.64 (2.05) | ns | ns |
| CREA (μmol/L) | 71.78 (15.36) | 68.22 (9.65) | 70 (12.69) | ns | ns |
| TBIL (μmol/L) | 14.64 (6.81) | 15.31 (7.41) | 14.9 (8.07) | ns | ns |
| UA (μmol/L) | 421.08 (143.1) | 376 (103.91) | 385.89 (104.27) | ns | ns |

**Table2.**
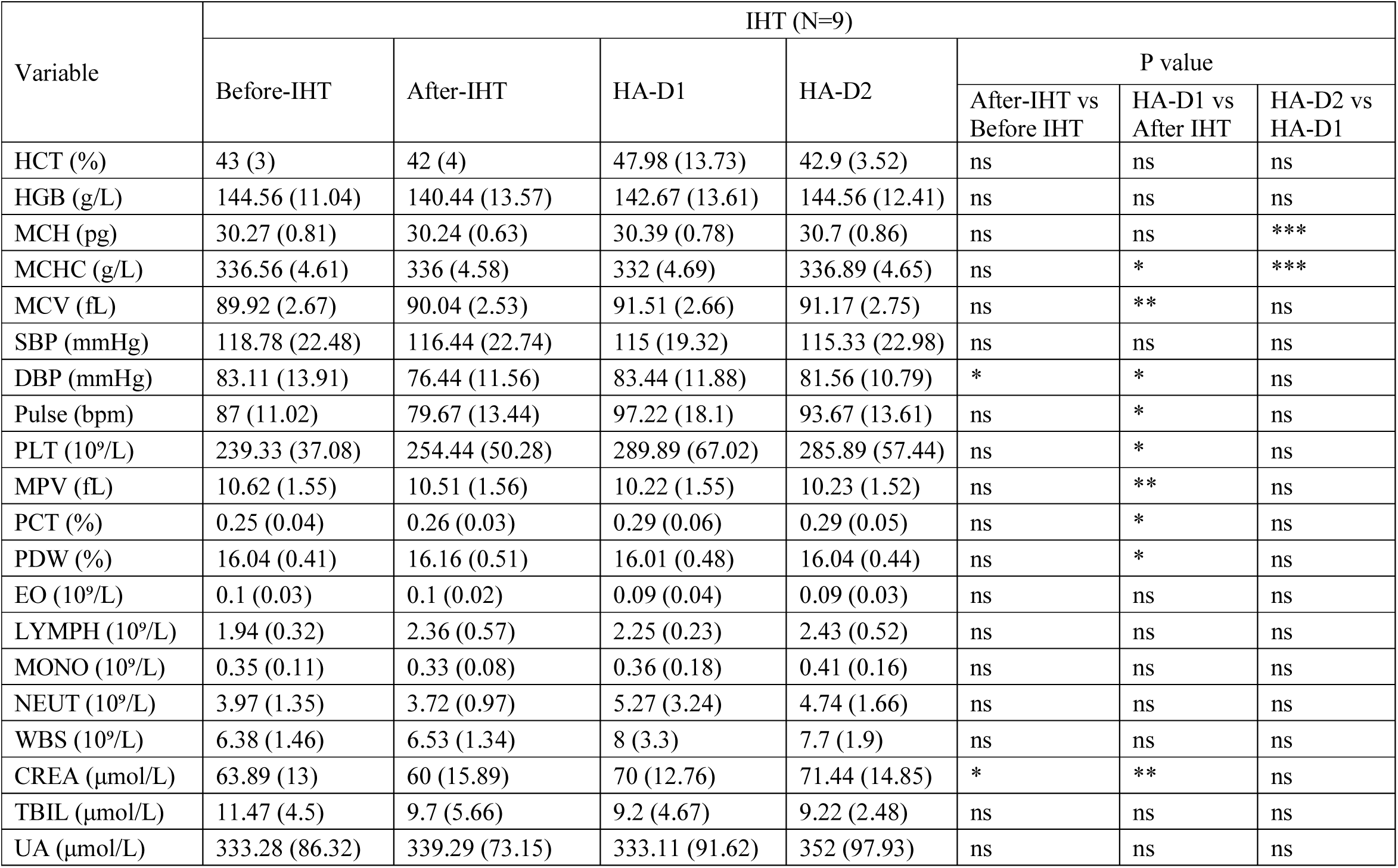
Demographic and clinical information of the IHT group.

Upon high-altitude exposure, the IHT group displayed significantly lower LLSs and higher peripheral oxygen saturation (SpO_2_) than the non-IHT control group **(Fig. 1B, C)**. No between-group differences in erythrocyte counts were observed, indicating that the protective effect of IHT was not attributable to increased erythropoiesis **(Fig. 1D)**. Consistent with this, hemoglobin (HGB) and hematocrit (HCT) levels showed no significant differences over the course of the experiment in either the control or IHT group **(Tables 1&2)**. These findings demonstrate that IHT induces rapid physiological adaptation to high-altitude hypoxia independent of changes in red blood cell number. Interestingly, mean corpuscular volume (MCV) decreased in both groups **(Tables 1&2)**, indicating that hypoxia influences erythrocyte volume regulation.

Given the abundance of mature erythrocytes (the most abundant cells in our body) and their established sensitivity to hypoxia, we performed untargeted metabolomics to characterize the systemic metabolic changes induced by IHT in both erythrocytes and plasma. We identified 300 metabolites in erythrocytes and 285 in plasma **(Supplemental Table1)**. In the control group, partial least-squares discriminant analysis (PLS-DA) of erythrocyte metabolites showed clear separation between sea-level and high-altitude time points. In contrast, in the IHT group, metabolic profiles after IHT (at sea level) and on HA-D1 were markedly more similar to each other than to the corresponding profiles in controls **(Fig. 2A, B)**. Plasma metabolomics revealed a comparable trend, albeit with less pronounced separation in the control group **(Fig. 2C, D)**. Hierarchical clustering further confirmed that metabolic profiles after IHT and on HA-D1 clustered more closely in the IHT group than in controls, both in erythrocytes and plasma **(Fig. 2A-D)**. Overall, these results indicate that IHT pre-configures the metabolic state of erythrocytes toward one that resembles initial acute high-altitude exposure in the control group, establishing a form of “hypoxic memory” that enhances subsequent hypoxic tolerance.

**Figure 2.**
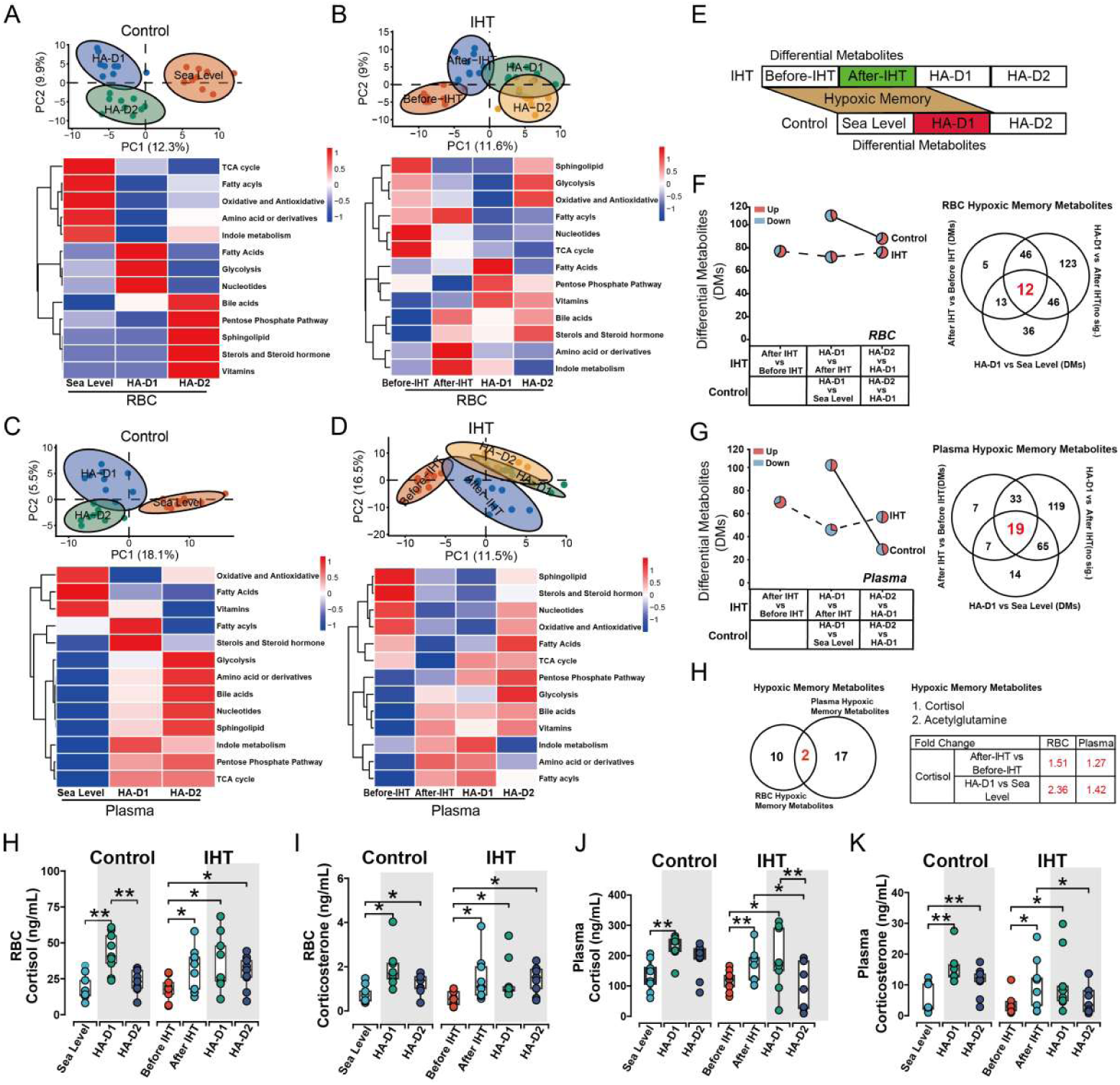
Intermittent Hypoxic Training Reprograms the Circulating Metabolome in Plasma and Red Blood Cells Prior to Altitude Exposure. (A-D) Partial least squares discriminant analysis (PLS-DA) and metabolic pathway enrichment. PLS-DA score plots display the distinct metabolomic profiles of red blood cells (RBCs; top panels, A, B) and plasma (top panels, C, D) from the Control (A, C) and IHT (B, D) groups. Ellipses represent 95% confidence intervals. Corresponding heatmaps below each score plot depict the abundance patterns of significantly altered metabolic pathways in the Control (A, RBCs; C, Plasma) and IHT (B, RBCs; D, Plasma) groups. **(E)** Schematic illustrating the proposed hypothesis of IHT-induced metabolic reprogramming and hypoxic memory formation. **(F)** Analysis of differentially abundant metabolites (DMs) in RBCs. The number of DMs identified across three key comparison pairs is shown: After-IHT vs. Before-IHT (within IHT group), HA-D1 vs. After-IHT (IHT group) or HA-D1 vs. Sea Level (Control group), and HA-D2 vs. HA-D1. The left panel uses pie charts to indicate the proportion of upregulated (red) and downregulated (blue) metabolites for each comparison. The right panel is a Venn diagram illustrating the overlap of unique and shared DMs from the various paired comparisons between the IHT and Control groups within RBCs. **(G)** Analysis of differentially abundant metabolites (DMs) in plasma. The layout mirrors panel F, showing the number, regulation direction (pie charts), and overlap (Venn diagram) of DMs identified in plasma across the same three comparison pairs. **(H)** Identification of hypoxic memory metabolites. The left panel is a Venn diagram illustrating the differentially abundant and shared “hypoxic memory” metabolites between RBCs and plasma. The right panel displays the fold changes of these metabolites across the various paired comparisons from the IHT and Control groups. **(I-L)** Targeted quantification of glucocorticoids via Multiple Reaction Monitoring (MRM). Cortisol levels were quantified in RBCs (I) and plasma (J) of Control and IHT groups. Corticosterone levels were quantified in RBCs (K) and plasma (L) of Control and IHT groups. Data are presented as mean ± SD. *p < 0.05, **p < 0.01 (paired two-tailed t-test).

To identify key metabolites involved in this adaptation, we performed differential abundance analysis for multiple comparisons, including After IHT vs. Before IHT (IHT group) and HA-D1 vs. Sea Level (control group). Stage-wise analysis revealed that most metabolic alterations in controls occurred during acute high-altitude exposure (HA-D1 vs. Sea Level), whereas in the IHT group, significant changes primarily followed the training intervention (After IHT vs. Before IHT) **(Fig. 2F,G)**. Notably, the IHT group exhibited far fewer differential metabolites during the acute transition to high altitude (HA-D1 vs. After IHT) than controls showed during their initial ascent (HA-D1 vs. Sea Level) **(Fig. 2F,G)**, indicating that IHT pre-adapts metabolism and attenuates additional metabolic reprograming upon HA exposure. Interestingly, in the control group, more metabolites were altered during the acute phase (HA-D1 vs. Sea Level) than during the subsequent day (HA-D2 vs. HA-D1), suggesting extensive compensatory adjustments following initial hypoxic stress **(Fig. 2F,G)**.

To pinpoint metabolites underlying this “hypoxic memory” **(Fig. 2E)**, we compared IHT-specific metabolites (i.e., those significantly changed after IHT but not further altered upon subsequent HA exposure) with those altered by acute high-altitude exposure in controls (HA-D1 vs. Sea Level). Venn diagram analysis identified 12 overlapping metabolites in erythrocytes and 19 in plasma (Fig. 1I). Cross-compartment analysis of these conserved “hypoxic memory” metabolites highlighted cortisol from the steroid hormone pathway as notably enriched in both compartments **(Fig. 2H)**. Targeted LC-MS/MS validation confirmed elevated cortisol levels in erythrocytes **(Fig. 2I, J)** and plasma **(Fig. 2K, L)** after IHT, whereas in controls, cortisol increased only after high-altitude exposure. A similar pattern was observed for corticosterone, indicating a broad glucocorticoid response to both IHT and acute hypoxia.

Collectively, these results demonstrate that IHT induces a metabolic state in erythrocytes and plasma, characterized by a marked increase in glucocorticoids, that resembles that observed following the initial acute hypoxic exposure in control individuals who had not received IHT. This metabolic preconditioning is associated with improved physiological adaptation to high altitude as judged by lower LLS on HA-D1. Our findings support a novel hypothesis that elevated glucocorticoids are key mediators of IHT-induced protection against high-altitude stress.

### Glucocorticoid Signaling via Glucocorticoid Receptors Enhances Erythrocyte Oxygen Release and Antioxidative Capacity by Inducing Glucose Metabolic Reprogramming

Glucocorticoids are well-established stress hormones known to mitigate inflammation and tissue damage under high-altitude stress. However, their potential role in mediating the “erythroid hypoxic metabolic memory” induced by IHT was unexplored prior to our research reported here.

Correlation analysis revealed that in individuals who underwent hypoxic preconditioning (After-IHT), glucocorticoid levels—whether in erythrocytes or plasma—were positively correlated with the post-ascent SpO₂ on high-altitude day 1 (HA-D1), suggesting a potential protective role. We separately examined the correlations of two glucocorticoids (cortisol and corticosterone) in erythrocytes and plasma with SpO₂ on HA-D1 (Figure 3A, B). The correlations were stronger in erythrocytes (cortisol: r = 0.85, p = 0.02; corticosterone: r = 0.75, p = 0.03) than in plasma (cortisol: r = 0.75, p = 0.02; corticosterone: r = 0.73, p = 0.01) (Figure 3A, B). These findings imply that after hypoxic preconditioning, glucocorticoid levels are closely linked to systemic oxygen saturation. Based on these observations, we proceeded to investigate the direct effects of glucocorticoids on erythrocyte function in vitro.

**Figure 3.**
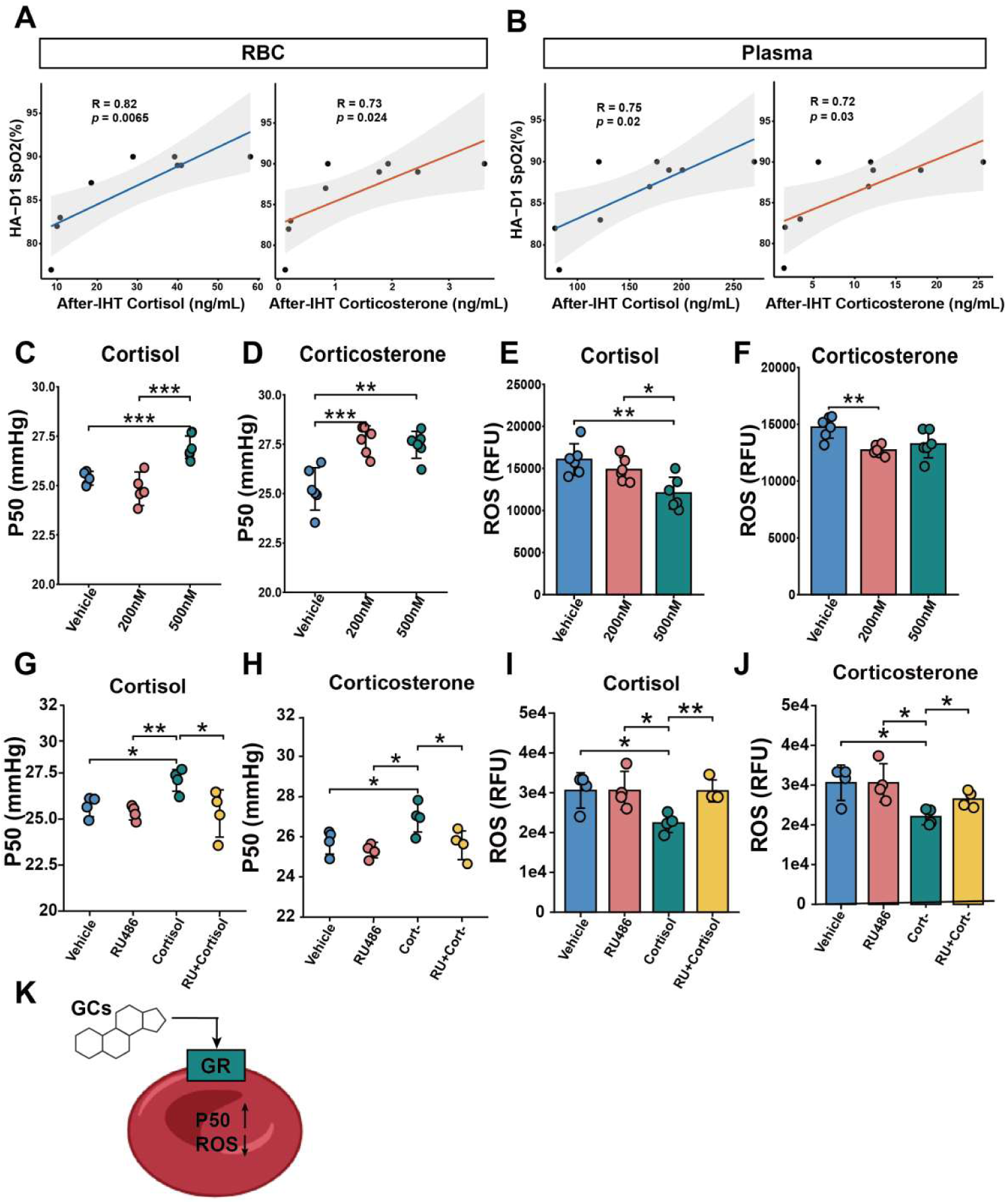
Glucocorticoids correlate with red blood cell physiology and modulate oxygen affinity and oxidative stress in vitro. **(A, B)** Correlations between glucocorticoid levels and Peripheral Oxygen Saturation (SpO_2_) in IHT group. Scatter plots show the associations between RBC and plasma cortisol or corticosterone levels with corresponding SpO_2_ upon high-altitude exposure on Day 1 (HA-D1). Pearson correlation coefficients (r) and corresponding p-values are indicated for each significant relationship. **(C-F)** In vitro modulation of RBC function by glucocorticoids. Purified human RBCs were treated with physiologically relevant concentrations of cortisol or corticosterone. The P50 value was measured following treatment with different concentration cortisol (C) or corticosterone (D). Intracellular ROS levels were quantified in RBCs treated with different concentration cortisol (E) or corticosterone (F). **(G-J)** In vitro modulation of RBC function by glucocorticoid in the presence of absence of a glucocorticoid receptor inhibitor. The P50 value was measured following treatment with cortisol (G) or corticosterone (H) Intracellular ROS levels were quantified in RBCs treated with cortisol (I) or corticosterone (J). Data are presented as mean ± SD of biologically independent replicates. *p < 0.05, **p < 0.01, ***p < 0.001 (one-way ANOVA with Dunnett’s post hoc test).**(K)** Schematic hypothesis illustrating the proposed regulatory role of glucocorticoids (GCs) in modulating RBC oxygen affinity (via P50, the partial pressure of oxygen at which hemoglobin is 50% saturated) and redox homeostasis (via reactive oxygen species, ROS).

Treatment of human erythrocytes with cortisol or corticosterone increased the P50 value (a measure of oxygen-release capacity) **(Fig. 3C, D)**. Concurrently, glucocorticoid treatment significantly reduced intracellular reactive oxygen species (ROS) levels **(Fig. 3E, F)**. To determine whether these effects were mediated via the glucocorticoid receptor (GR), we co-cultured erythrocytes with glucocorticoids in the presence or absence of the specific GR antagonist RU486. The results show that RU486 completely abolished the glucocorticoid-induced increase in P50 and decrease in ROS, confirming that GR signaling is essential for these functional adaptations **(Fig. 3G-J).**

Since glucose metabolic reprogramming is central to erythrocyte adaptation under hypoxia, we next asked whether glucocorticoids enhance erythrocyte function by modulating glucose metabolism **(Fig. 4).** We performed ^13^C-glucose metabolic tracer analysis using erythrocytes treated with or without glucocorticoids, and in the presence or absence of the GR antagonist RU486. Glucocorticoid exposure rapidly accelerated glycolysis within 30 minutes, as evidenced by increased uptake of glucose(M+6) and elevated production of lactate(M+3). Key glycolytic metabolites, including the critical oxygen-affinity modulator 2,3-bisphosphoglycerate (2,3-BPG), were also significantly elevated **(Fig. 4)**. Concurrently, intermediates of the pentose phosphate pathway (PPP), such as 6-phosphogluconolactone (6PGL), 6-phosphogluconate (6PG), ribose-5-phosphate (R5P), sedoheptulose-7-phosphate (S7P), and erythrose-4-phosphate (E4P), showed marked increases upon glucocorticoid treatment **(Fig. 4)**. Notably, these glucocorticoid-induced metabolic shifts were largely abolished by co-treatment with RU486, indicating GR mediated signaling. Compared to the controls, erythrocytes treated with glucocorticoids plus RU486 exhibited significantly lower levels of glycolytic and pentose phosphate pathway intermediates than those treated with glucocorticoids alone **(Fig. 4),** confirming that GR activation mediates the metabolic reprogramming triggered by glucocorticoids.

**Figure 4.**
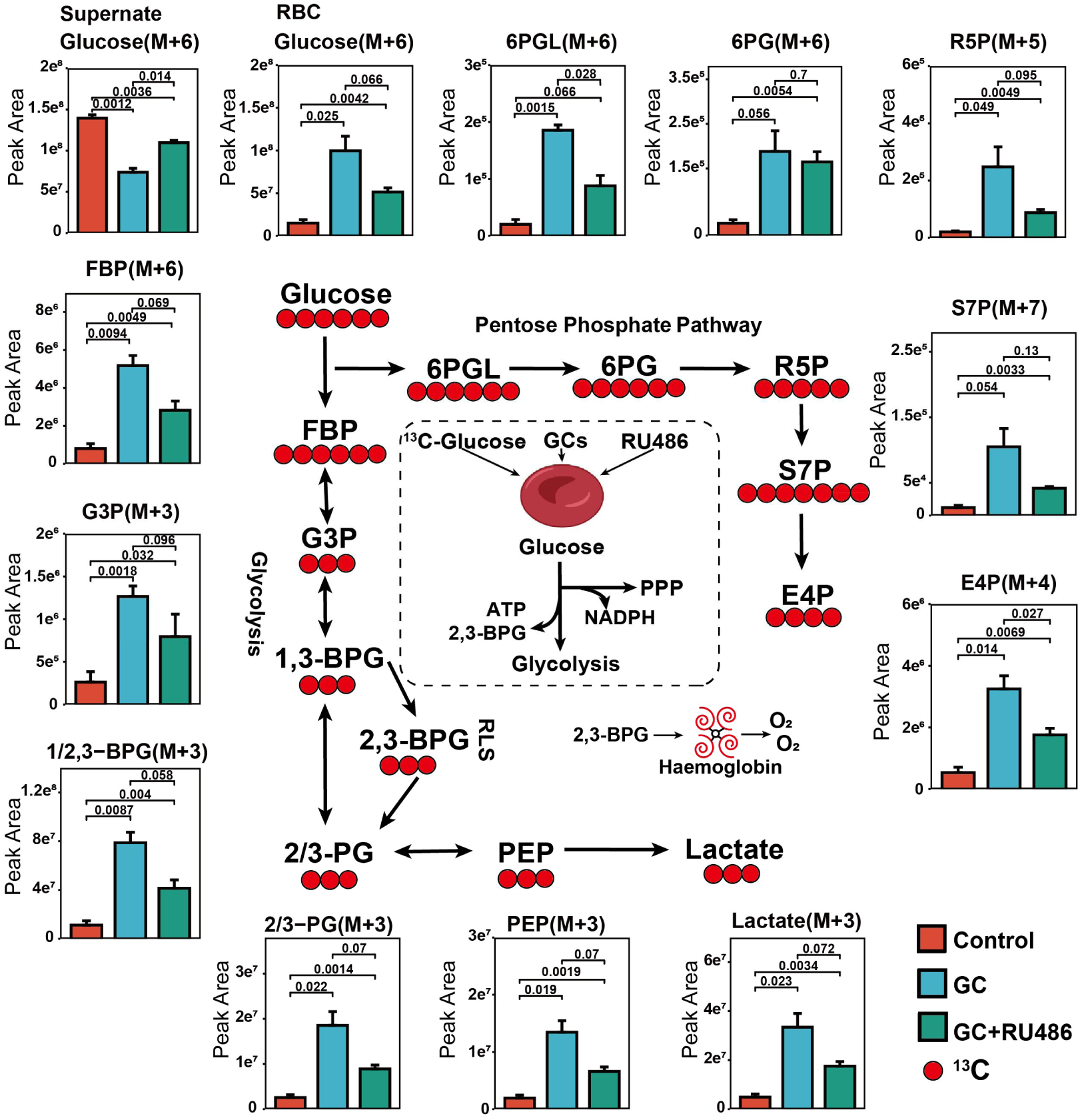
Glucocorticoids reprogram human red blood cell energy metabolism in vitro by enhancing glycolysis and the pentose phosphate pathway. Schematic of central glucose metabolism pathways in RBCs, glycolysis and the pentose phosphate pathway. Key measured metabolites and their ¹³C-labeled isotopologues are indicated along the pathways. Bar plots showing the peak areas of ¹³C-labeled metabolites in the supernatant (cell culture medium) and within red blood cells (RBCs) under different treatment conditions: vehicle control (Control, red), glucocorticoid treatment (GC, blue), and co-treatment with glucocorticoid and the glucocorticoid receptor antagonist RU486 (GC+RU486, green). Peak Area are presented as mean ± SD of biological replicates. The exact p-values for significant comparisons are annotated above the bars (unpaired two-tailed t-test for pairwise comparisons between indicated groups).

Collectively, these data demonstrate that glucocorticoids, acting through GRs, rapidly reprogram erythrocyte glucose metabolism toward enhanced glycolysis and pentose phosphate pathway activity. This metabolic reprogramming increases 2,3-BPG synthesis, thereby facilitating oxygen offloading, and likely bolsters antioxidative capacity via PPP-mediated NADPH production, providing a mechanistic basis for the observed improvements in P50 and ROS clearance **(Fig. 4)**.

### IHT Remodels the Erythrocyte Lipidome and Activates a Glucocorticoid-Driven Sphingolipid Axis

In view of the established link between erythrocyte lipid metabolism and systemic oxygen delivery, we performed untargeted lipidomics to determine if lipid metabolism is altered by IHT. The results identified 789 and 809 lipid species in erythrocytes and plasma, respectively **(Supplemental Table3)**. Partial least squares-discriminant analysis (PLS-DA) revealed distinct clustering in different groups. In controls, erythrocyte lipidomes clearly separated across sea level (SL), HA-D1, and HA-D2, whereas in the IHT group, post-training samples clustered closely with those from high-altitude exposure (HA-D1, HA-D2) of the control group **(Fig. 5A)**. A similar shift was observed in plasma **(Fig. 5B)**. These findings indicate that IHT remodels the lipid profile toward a state resembling HA-D1 of the control group.

**Figure 5.**
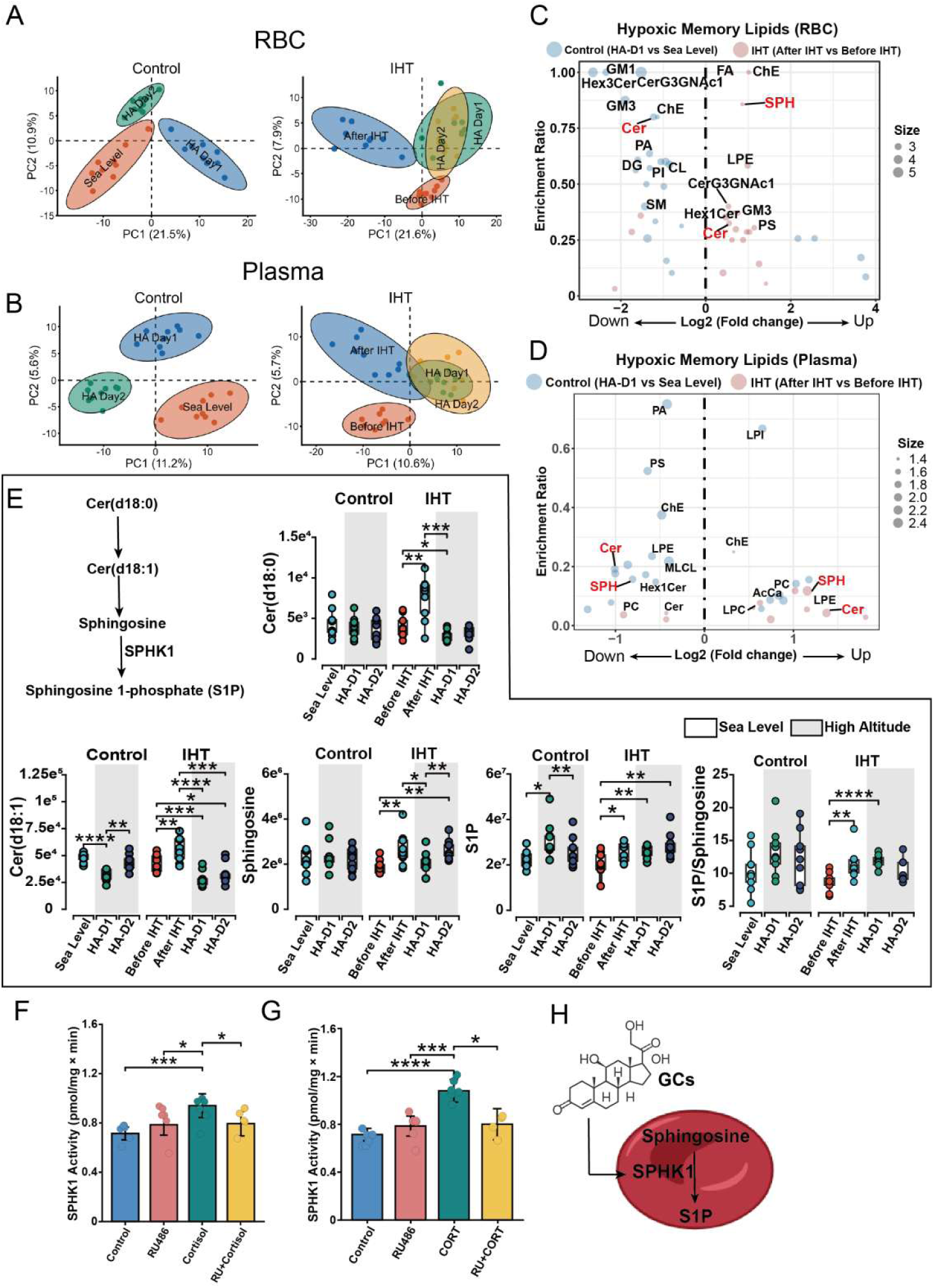
Lipidomic profiling reveals a specific enhancement of the ceramide metabolic pathway in RBCs following intermittent hypoxic training. **(A, B)** Partial least squares discriminant analysis (PLS-DA) score plots display the global lipid profiles of RBCs (A) and plasma (B) from Control and IHT groups. Ellipses represent 95% confidence intervals. **(C, D)** Valcano plot of lipid class reveals the key lipid class alterations After-IHT vs Before-IHT in IHT group (red) and HA-D1 vs Sea Level in control group (blue). Corresponding dot show the enrichment ratios of significantly altered lipid metabolic pathways in RBCs (C) and plasma (D). **(E)** Schematic of the ceramide metabolism pathway and the abundance of ceramide pathway intermediates. Box plots show the relative levels of key metabolites (ceramides, sphingosine, S1P) within the ceramide metabolic pathway in RBCs of Control and IHT groups. Data are mean ± SD. *p < 0.05, **p < 0.01, ***p < 0.001, ****p < 0.0001 (paired two-tailed t-test). **(F)** The diagram highlights the key enzyme sphingosine kinase-1 (SPHK1). **(G, H)** Enzymatic activity of SPHK1 in purified RBCs treated in vitro with vehicle, glucocorticoid (GC) or GC+ glucocorticoid receptor inhibitor. Data are mean ± SD. *p < 0.05, **p < 0.01 (unpaired two-tailed t-test (G) or one-way ANOVA with Dunnett’s test (H)).

Volcano plot analysis comparing lipids altered post-IHT (After IHT vs. Before IHT) with those altered post-altitude in controls (HA-D1 vs. SeaLevel) showed that most differential lipids were upregulated after IHT but downregulated after altitude exposure in controls. Ceramides (Cer) were upregulated in erythrocytes after IHT and downregulated in controls, a pattern mirrored in plasma **(Fig. 5C&D)**. We therefore hypothesized that ceramides contribute to IHT-induced adaptation. Given that ceramide is a precursor of sphingosine-1-phosphate (S1P) **(Fig. 5E)**, a lipid mediator involved in hypoxic responses, we proposed that IHT augments ceramide and sphingosine metabolism to establish a metabolic reserve for subsequent hypoxic stress. Specific ceramide species-Cer(d18:0) and Cer(d18:1)-along with sphingosine and S1P, were elevated after IHT, whereas in controls, S1P rose only upon high-altitude exposure **(Fig. 5E)**.

In vitro, glucocorticoids potently activated erythrocyte SPHK1, an effect blocked by the GR antagonist RU486, identifying glucocorticoid-GR signaling as an upstream regulator of SPHK1 activity **(Fig. 5F-H)**. Collectively, these data suggest that IHT increases ceramide availability and enhances SPHK1 activity via glucocorticoid signaling to boost S1P production in erythrocytes.

### Erythrocyte SPHK1 Functioning Downstream of Glucocorticoid Receptor Activation Counteracts Hypoxia

To determine whether elevated glucocorticoids can precondition erythrocyte function in vivo and confer systemic protection, we performed preclinical studies in mice. Mice were pretreated with glucocorticoids (cortisol and corticosterone) for three days prior to hypoxic exposure **(Fig. 6A)**. Under subsequent hypoxia, glucocorticoid-pretreated mice exhibited significantly higher erythrocyte P50 and markedly lower intracellular ROS levels compared to untreated hypoxic controls **(Fig. 6D&E)**. Consistent with improved systemic oxygen delivery, renal hypoxia, assessed via a hypoxia-sensitive fluorescent probe, was substantially attenuated in glucocorticoid pretreated mice exposed to hypoxia, maintaining low levels comparable to normoxic controls **(Fig. 6F&G)**.

**Figure 6.**
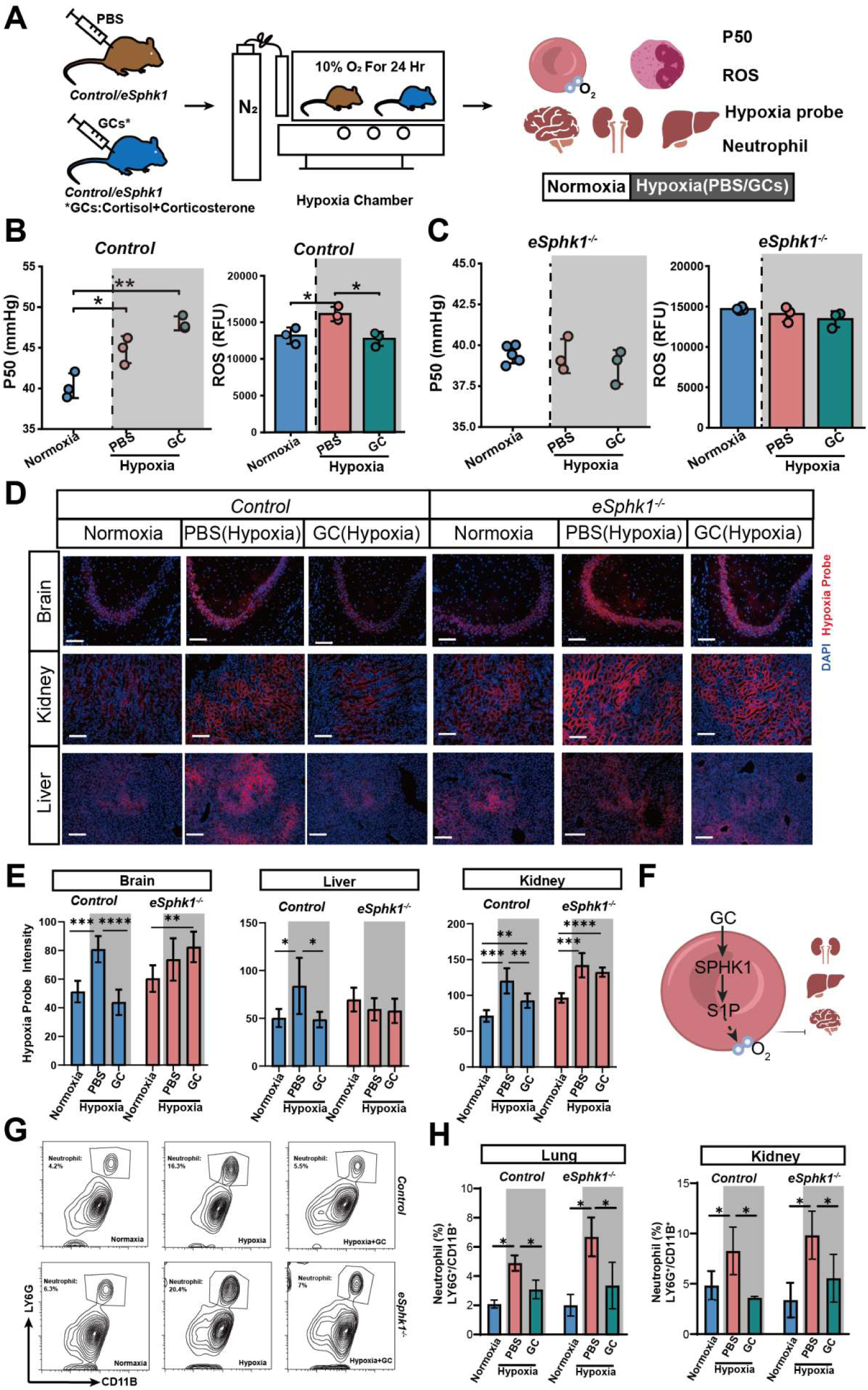
Glucocorticoids induce SPHK1 and enhance oxygen release to alleviate organ hypoxia. **(A)** Experimental schematic. Mice were pre-treated with glucocorticoid (GC) or PBS for 3 days, followed by exposure to normoxia or hypoxia (10% O₂ for 24 hr). **(B, C)** SPHK1 mediates GC-induced changes in oxygen affinity and oxidative stress. The GC-induced rightward shift of the hemoglobin oxygen dissociation curve (increased P50) and the reduction in intracellular ROS levels in erythrocytes (B) are abrogated in *eSphk1^-/-^*mice (C). **(D)** In vivo imaging of tissue hypoxia. Fluorescence images of brain, kidney, and liver stained with a hypoxia probe (red) and DAPI (blue nuclear stain) from control and *eSphk1^-/-^* mice under the indicated conditions. Scale bar = 50 μm. **(E)** Quantification of tissue hypoxia. Quantification of hypoxia probe intensity in brain, kidney, and liver. Data are mean ± SEM. *p < 0.05, **p < 0.01, ***p < 0.001, ****p < 0.0001 (two-way ANOVA with Tukey’s post hoc test). **(F)** Schematic model of the proposed “GC-GR-SPHK1” axis. Glucocorticoids (GCs) activate SPHK1 signaling in red blood cells, which enhances their oxygen-release capacity, thereby alleviating hypoxia in peripheral tissues. **(G, H)** GC-induced neutrophilia is attenuated in *eSphk1^-/-^* mice. Quantification of neutrophil infiltration (LY6G^+^/CD11B^+^) in lung (B) and kidney (C) tissues following the treatments described in (A).

To define the specific contribution of erythrocyte SPHK1 to this glucocorticoid-mediated adaptation, we utilized erythrocyte specific *Sphk1*-knockout mice (*eSphk1^-/-^*). In contrast to wild-type littermates **(Fig. 6D)**, *eSphk1^-/-^* mice showed no improvement in erythrocyte P50 or reduction in ROS following glucocorticoid pretreatment and subsequent hypoxia **(Fig. 6E)**. Assessment of hypoxia in the kidney, brain, and liver using hypoxia-sensitive probes revealed that glucocorticoid pretreatment markedly attenuated tissue hypoxia in control mice exposed to hypoxia. In contrast, this protective effect was completely abolished in *eSphk1^-/-^* mice, in which hypoxia levels in these organs remained comparable to those in untreated, hypoxic controls **(Fig. 6F&G)**. These results establish that erythrocyte SPHK1 is essential for mediating the glucocorticoid-induced enhancement of systemic oxygen delivery in vivo. Together, these findings demonstrate that elevated glucocorticoids activate an erythrocyte-intrinsic “GC-GR-SPHK1-S1P” axis, which is indispensable for enhancing systemic hypoxia tolerance following intermittent hypoxic training **(Fig. 6H)**

To assess the systemic immunomodulatory effect of glucocorticoid pretreatment to subsequent hypoxia challenge, we measured neutrophil populations in the lungs and kidneys by flow cytometry. Glucocorticoid pretreated mice exhibited significantly fewer neutrophils in both tissues after hypoxia compared to untreated hypoxic controls **(Fig. 6B, C)**, indicating that glucocorticoid pretreatment attenuates hypoxia-induced neutrophilic inflammation. Notably, this reduction in neutrophil infiltration was preserved in *eSphk1^-/-^* mice, demonstrating that the systemic anti-inflammatory effect of glucocorticoids is independent of erythrocyte-intrinsic SPHK1 signaling **(Fig. 6B, C)**.

These results establish that erythrocyte SPHK1 is essential for mediating the glucocorticoid-induced enhancement of systemic oxygen delivery in vivo. Together, these findings demonstrate that elevated glucocorticoids activate an erythrocyte-intrinsic “GC-GR-SPHK1-S1P” axis, which is indispensable for enhancing systemic hypoxia tolerance following intermittent hypoxic training **(Fig. 6H).**

## DISCUSSION

This study fundamentally redefines the therapeutic paradigm of glucocorticoids in high-altitude medicine. Rather than merely treating AMS through established anti-inflammatory effects, we show that glucocorticoids function as endogenous IHT-induced preconditioning agents that stimulate erythrocyte hypoxic memory before ascent. This represents a conceptual inversion from “rescue therapy” to metabolic pre-conditioning. Our discovery of the “GC-GR-SPHK1-S1P” axis establishes the first non-transcriptional mechanism by which glucocorticoids operate in terminally differentiated erythrocytes. By coupling enhanced glucose metabolism and sphingolipid signaling, the “GC-GR-SPHK1-S1P” signaling pathway transforms erythrocytes from passive oxygen carriers into adaptive metabolic sensors that preemptively optimize oxygen delivery. Clinically, we establish erythrocyte metabolic plasticity as a druggable target and glucocorticoids as rapid-acting conditioning hormones, opening new avenues for preventing hypoxic end-organ injury across diverse hypoxia-driven clinical contexts (**Fig.7**).

Our study adds a transformative chapter to glucocorticoid biology. Beyond their canonical role as potent anti-inflammatory agents to treat AMS^35–37^, they function as rapid-acting metabolic hormones that stimulate erythrocyte hypoxic memory—converting passive oxygen carriers into adaptive, self-tuning oxygen delivery vehicles primed for ascent. Notably, in humans we unexpectedly found that IHT induces both plasma and erythrocyte glucocorticoid levels and showed a negative correlation between pre-ascent glucocorticoid levels and subsequent high altitude AMS severity. This finding prompted us to further dissect the protective function of glucocorticoids and underlying mechanism in erythrocytes. Functionally, we provide both in vitro and preclinical in vivo evidence that glucocorticoid treatment induces oxygen delivery and anti-ROS capacity in cultured primary human erythrocytes and in erythrocytes from wild type mice. Significantly, the glucocorticoid treatment attenuated tissue hypoxia and pulmonary neutrophil infiltration in vivo.

We employed metabolomics and isotopically labelled glucose tracer analyses to demonstrate that glucocorticoid signaling via GR activation enhanced glucose uptake and its metabolism through glycolysis and the pentose phosphate pathway (PPP). Increased glycolysis was accompanied by increased metabolism through the Rapoport-Luebering shunt (RLS), resulting in a marked increase in the synthesis of 2,3-bisphosphoglycerate (2,3-BPG), a critical allosteric effector that reduces hemoglobin-oxygen affinity, directly enhancing the oxygen-release capacity of erythrocytes (increased P50)^38^. Increased metabolism through the PPP results in increased production of NADPH, thereby providing enhanced anti-oxidative ability to protect erythrocytes from ROS. Thus, we provide a previously unrecognized protective “hypoxic metabolic memory” resulting from glucocorticoid-mediated induction of erythrocyte 2,3-BPG production to promote enhanced oxygen delivery as well as increased metabolism through the PPP for better anti-oxidative capacity to combat tissue hypoxia upon ascent.

Our lipidomics analysis revealed that IHT induces sphingolipid metabolic reprograming toward S1P production, with S1P levels positively correlating with glucocorticoid levels in humans. This discovery enabled us to identify SPHK1 as a critical GR downstream effector in glucocorticoid signaling. Genetic proof-of-principle studies in *eSphk1^-/-^*mice demonstrated that removal of erythrocyte SPHK1 abolishes glucocorticoid-mediated hypoxia protection, resulting in defective oxygen delivery, severe tissue hypoxia, and exaggerated neutrophil infiltration. These findings align with prior evidence that S1P stimulates increased glycolysis by promoting the release of GAPDH from membrane sequestration and stimulates increased 2,3-BPG production by promoting the phosphorylation and activation of 2,3-BPG mutase^23^. Collectively, we provide significant new insight that GR signaling enhances erythrocyte oxygen delivery and anti-oxidative resilience through the activation of SPHK1 and increased S1P synthesis and anti-oxidative resilience through increase metabolism through the PPP.

Overall, our study redefines erythrocytes as hormone-responsive hypoxic memory units and glucocorticoids as adaptive preconditioning signals, establishing the “GC-GR-SPHK1-S1P” axis as a druggable signaling pathway for rapid hypoxic tolerance. By targeting this axis, “IHT-mimetic” pharmacology can bypass weeks of preconditioning to deliver immediate metabolic resilience, based on glucocorticoid mediated oxygen delivery optimization. In addition to high altitude acclimatization this paradigm offers a potential universal strategy for inducible hypoxic memory in at-risk populations including those with hypoxia-related peripheral disease.

## Nonstandard Abbreviations and Acronyms

AMS: acute mountain sickness
IHT: Intermittent hypoxia training
2,3-BPG: 2,3-bisphosphoglycerate
GC: glucocorticoid
LLS: Lake Louise Score
Cer: Ceramides
S1P: sphingosine-1-phosphate
GR: glucocorticoid receptor
SPHK1: sphingosine kinase 1

## Acknowledgments

We deeply appreciate the dedication and commitment of the volunteers who participated in this study. All acknowledged individuals provided written consent (via email) for their inclusion in this section.

---

**Supplemental Figure 1.**
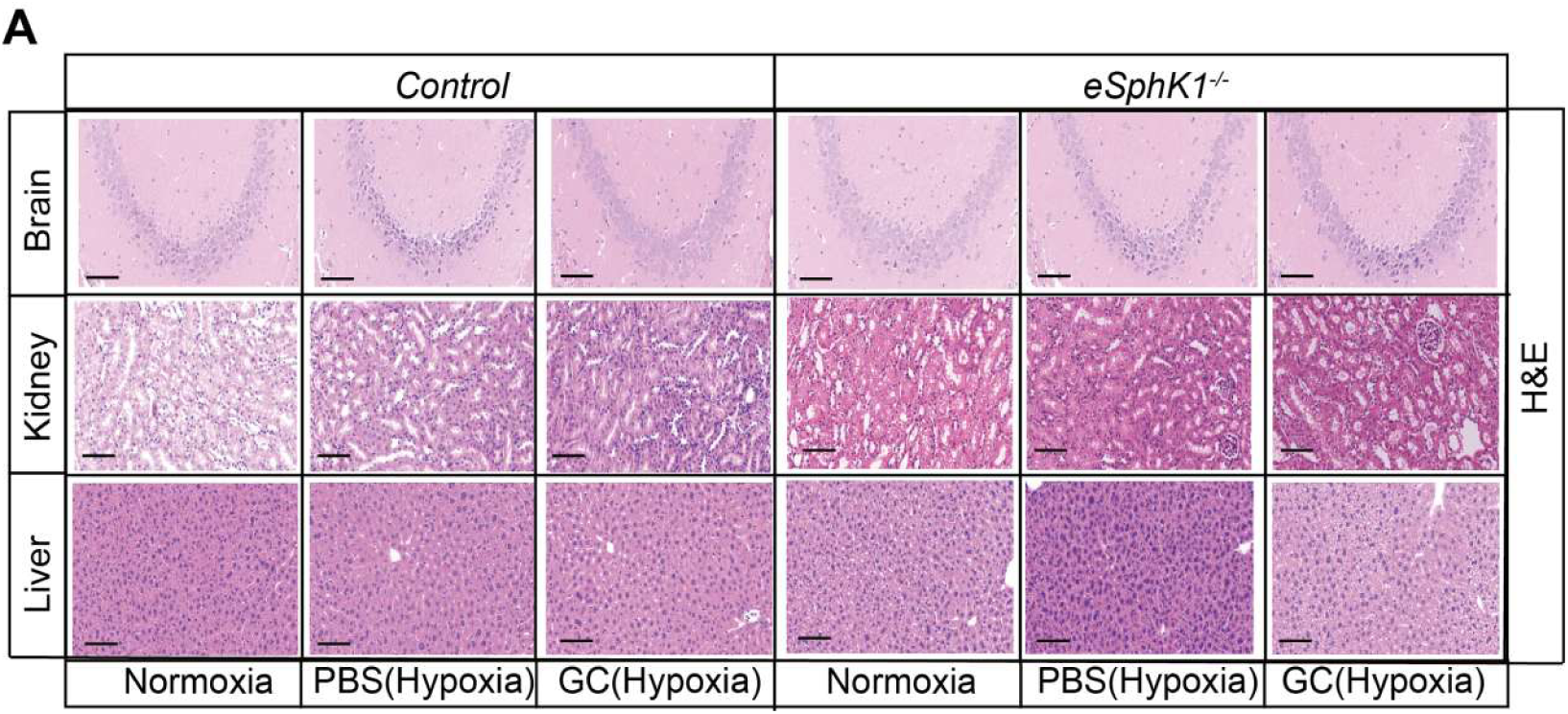
Histopathological assessment of brain, kidney, and liver tissues. Representative hematoxylin and eosin (H&E)-stained sections from the cerebral cortex (hippocampal region), kidney, and liver of control and *eSphk1^-/-^* mice.

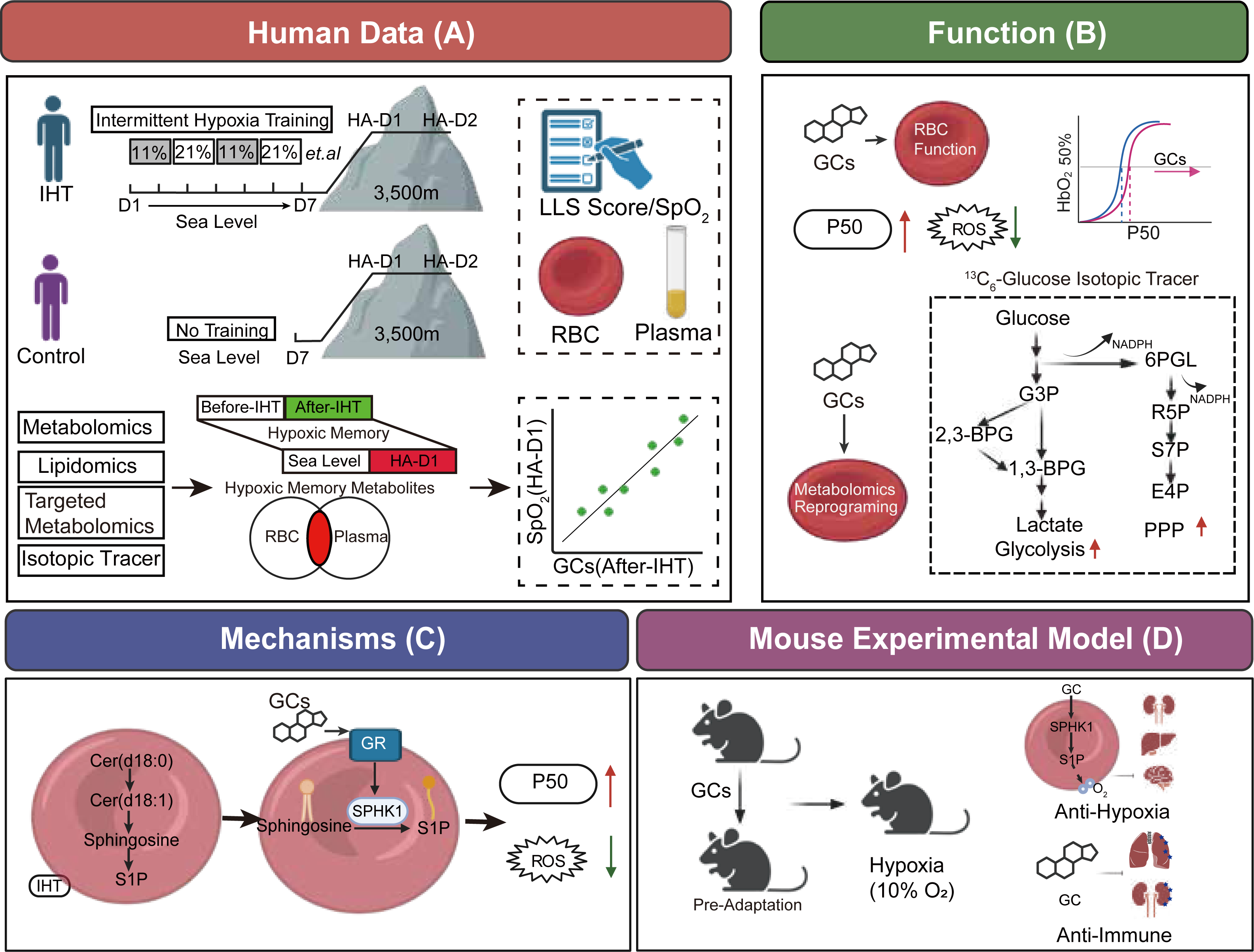

